# ImmuCellAI: a unique method for comprehensive T-cell subsets abundance prediction and its application in cancer immunotherapy

**DOI:** 10.1101/872184

**Authors:** Ya-Ru Miao, Qiong Zhang, Qian Lei, Mei Luo, Gui-Yan Xie, Hongxiang Wang, An-Yuan Guo

## Abstract

The distribution and abundance of immune cells, particularly T-cell subsets, play pivotal roles in cancer immunology and therapy. There are many T-cell subsets with specific function, however current methods are limited in estimating them, thus, a method for predicting comprehensive T-cell subsets is urgently needed in cancer immunology research. Here we introduce Immune Cell Abundance Identifier (ImmuCellAI), a novel gene set signature-based method, for precisely estimating the abundance of 24 immune cell types including 18 T-cell subsets, from gene expression data. Performance evaluation on both our sequencing data with flow cytometry results and public expression data indicated that ImmuCellAI can estimate immune cells with superior accuracy than other methods especially on many T-cell subsets. Application of ImmuCellAI to immunotherapy datasets revealed that the abundance of dendritic cells (DC), cytotoxic T, and gamma delta T cells was significantly higher both in comparisons of on-treatment vs. pre-treatment and responders vs. non-responders. Meanwhile, we built an ImmuCellAI result-based model for predicting the immunotherapy response with high accuracy (AUC 0.80~0.91). These results demonstrated the powerful and unique function of ImmuCellAI in tumor immune infiltration estimation and immunotherapy response prediction. The ImmuCellAI online server is freely available at http://bioinfo.life.hust.edu.cn/web/ImmuCellAI/.

## 1. Introduction

The immune system, comprising various proteins, immune cells, and tissues, is complex and important for host defense^[1]^. Immune cells, including innate immune cells [e.g., macrophages, neutrophils, natural killer (NK) cells, and dendritic cells (DC)] and adaptive immune cells (e.g., B and T cells), are important components of the immune system. Dysfunctions of immune cells such as abnormal distributions with respect to abundance and type as well as abnormal development and functions are always associated with diseases, including cancers^[2,3]^. Thus, investigating immune cell distribution in individuals could provide important insights into immune status, disease progression and prognosis, and therapy (particularly in cancer immunotherapy)^[4]^.

Tumor-infiltrating immune cells are considered to be primary immune signatures and are strongly associated with the clinical outcomes of immunotherapies^[5]^. T cells play pivotal roles in cancer initiation, progression, and therapy (particularly immunotherapy)^[6]^ and are composed of two major groups, each including numerous functional subpopulations (or subsets): CD4^+^ and CD8^+^ populations. The CD4^+^ T-cell subsets, such as T helper cells (e.g., Th1, Th2, Th17, and Tfh) and regulatory T cells (e.g., nTreg, iTreg, and Tr1), primarily display helper and/or regulatory activities on other immune cells^[7]^. The CD8^+^ T-cell subsets, cytotoxic T cells (Tc) and mucosal-associated invariant T cells (MAIT), function in killing target cells. Importantly, the abundance of T-cell subsets, particularly that of tumor-infiltrating T cells, could influence clinical curative effects and prognosis^[8]^. In addition, strategies used for regulating the proportion of T-cell subsets have demonstrated profound efficacy in cancer immunotherapies. For example, increasing the ratio of Teff/Treg subsets could enhance the antitumor effects of anti-CTLA-4 therapy against melanoma^[9]^. Thus, investigating the landscape of immune cells, particularly T cells, can help us better understand the interplay between the immune system and diseases and provide important clues for improving the efficacy of immunotherapy in precision medicine^[10]^.

High-throughput technologies, including microarrays and RNA sequencing (RNA-Seq), produce large-scale transcriptome data and provide opportunities for estimating the abundance of immune cells using gene expression profiles. Several methods, including xCell^[11]^, CIBERSORT^[12]^, EPIC^[13]^, TIMER^[14]^, and MCP-counter^[15]^, have been developed for enumerating immune cells from bulk transcriptome data of tumor samples, whereas rare method has been designed for estimating the abundance of numerous T-cell subsets, such as iTreg, Tc, and exhausted T cells (Tex). As such, there is an urgent need to develop a method focusing on abundance prediction of T-cell subsets and other important immune cells in immuno-oncology and immunotherapy studies.

In this study, we developed Immune Cell Abundance Identifier (ImmuCellAI), a method to robustly and precisely estimate the abundance of 24 immune cell types (including 18 T-cell subsets) from transcriptome data. ImmuCellAI was suitable for application to both microarray expression and RNA-Seq data from various resources (e.g., tumor, adjacent or normal tissue, and peripheral blood). Furthermore, we applied ImmuCellAI to cancer immunotherapy and The Cancer Genome Atlas (TCGA) pan-cancer data to explore the influence of immune cells on the efficacy of immunotherapy and clinical progression of patients with cancer.

## 2. Results

### 2.1. Algorithmic overview of the ImmuCellAI method

ImmuCellAI was designed to estimate the abundance of 18 T-cell subsets [CD4^+^, CD8^+^, CD4^+^ naïve, CD8^+^ naïve, central memory T (Tcm), effector memory T (Tem), Tr1, iTreg, nTreg, Th1, Th2, Th17, Tfh, Tc, MAIT, Tex, gamma delta T (γδ T), and natural killer T (NKT) cells] and six other important immune cells [B cells, macrophages, monocytes, neutrophils, DC, and NK cells] (Figure 1A). A brief illustration of the core algorithm of ImmuCellAI is represented in Figure 1B, and its detailed algorithm is described in the Online Methods section. Briefly, we curated a specific gene set as a gene signature for each immune cell type (table S1) from published reports and obtained its reference expression profile from the Gene Expression Omnibus (GEO) database (table S2). Then, we calculated the total expression deviation of the gene signature in the input expression dataset in comparison with the reference expression profiles of the 24 immune cell types. We assigned the deviation to the corresponding immune cell type based on the enrichment score of its gene signature, which was calculated using the single sample gene set enrichment analysis (ssGSEA) algorithm^[16]^. To correct the bias due to shared genes in the gene signatures of different immune cell types, a compensation matrix and least square regression were implemented to measure the weight of shared genes on these immune cells and to re-estimate their abundance (Figure 1B). ImmuCellAI was suitable for application to both RNA-Seq and microarray expression data from blood or tissue samples. To better utilize ImmuCellAI, we designed a user-friendly web server, which is freely available at http://bioinfo.life.hust.edu.cn/web/ImmuCellAI/, for estimating the abundance of 24 immune cell types from gene expression profiles.

**Figure1.**
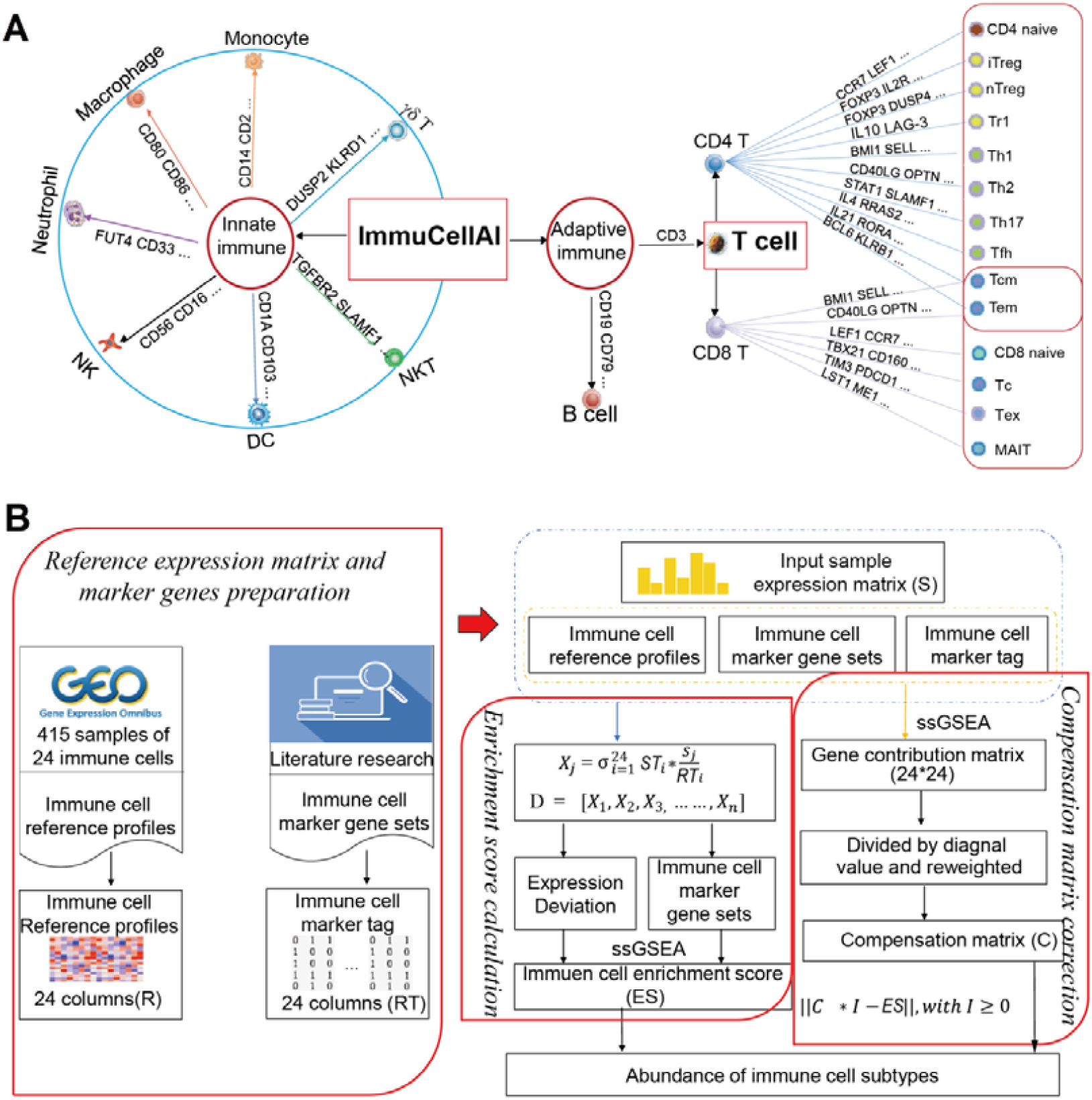
Immune cell types estimated by ImmuCellAI and the workflow of ImmuCellAI. **A.** Immune cell subsets enumerated by ImmuCellAI. Genes on the line to cell types are the examples of their marker genes. B. The pipeline of the ImmuCellAI algorithm. The three red boxes are the three main steps of ImmuCellAI algorithm. The reference expression profiles of the immune cells were obtained from GEO, and marker genes per immune cell type were obtained from the literature and analytical methods. For each queried sample, the enrichment score of total expression deviation of the signal gene sets was calculated and assigned to each immune cell type by the ssGSEA algorithm. The compensation matrix and least square regression were implemented to correct the bias caused by the shared marker genes among different immune cell types.

### 2.2. Performance of ImmuCellAI in RNA-Seq and microarray datasets

To evaluate the performance of ImmuCellAI, we applied it to multiple RNA-Seq and microarray expression datasets, performed benchmark tests, and compared the results using five other methods (xCell^[11]^, CIBERSORT^[12]^, EPIC^[13]^, MCP-counter^[15]^, and TIMER^[14]^). The Pearson correlation between the abundance estimated by flow cytometry and *in silico* method was used to assess the performance of each method in estimating the abundance of individual immune cell type, whereas the correlation deviation for all cell types was calculated to systematically evaluate the overall prediction power of each method (details are discussed in the Online Methods section).

First, we enumerated the amount of immune cell types using all six analytical methods, among which ImmuCellAI proved capable of predicting more T cell subsets than the other methods (Figure 2A). Then, we used six RNA-Seq datasets as benchmark resources for evaluating the performance of ImmuCellAI (Figure 2B and 2C). Each dataset contained counting results of immune cells identified by flow cytometry of samples. Three of them were simulated and integrated from single-cell sequencing data of liver cancer (GSE98638)^[18]^, lung cancer (GSE99254)^[19]^, and melanoma (GSE72056)^[20]^. One dataset was taken from the lymph nodes of four patients with melanoma included in the EPIC^[13]^ project. Furthermore, because of the limited number of T-cell subsets in currently available data, to evaluate the performance of ImmuCellAI in estimating the abundance of unique T-cell subsets, we generated two datasets using flow cytometry analysis for all 24 immune cell types and sequenced their RNA (BIGD id: CRA001839 and CRA001840). One of these datasets contained five samples from healthy donors and the other contained seven samples from patients with acute myelocytic leukemia. Based on the results, the abundance of most immune cells estimated by ImmuCellAI showed a higher positive correlation with the counting results of flow cytometry than that estimated by the other methods, particularly for T-cell subsets. These results suggested that ImmuCellAI robustly and accurately enumerates the 24 immune cell types in RNA-Seq datasets (Figure 2B and 2C; Figure S1A, S1B and S2A).

**Figure2.**
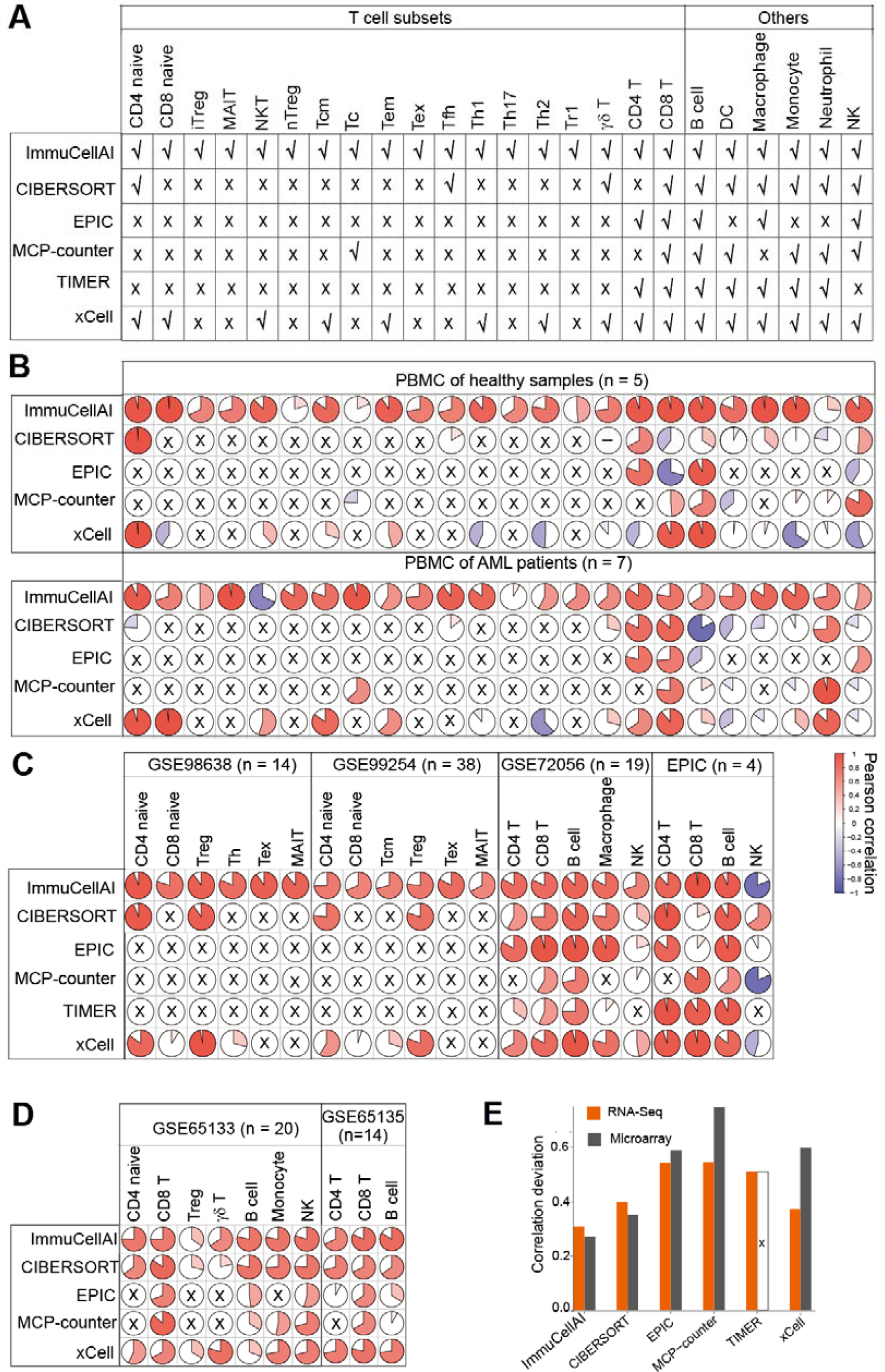
Performance comparison of ImmuCellAI and other methods. **A.** Immune cell types can be estimated in ImmuCellAI and other five methods (CIBERSORT, EPIC, MCP-counter, TIMER, and xCell). B. Prediction accuracy of ImmuCellAI and other methods for our sequenced blood samples from healthy individuals and AML patients. The rows correspond to methods and the columns indicate the Pearson coefficient for the corresponding cell in the pie graph. Cell types not available in the corresponding methods are marked with a black “×.” The “–” in the circle denotes the correlation analysis result was “NA.” **C.** Performance of ImmuCellAI and other methods when applied to public RNA-Seq datasets. D. Performance of ImmuCellAI and other methods on microarray datasets. E. Correlation deviation of each method, which took sample size and overall accuracy into consideration to measure the global performance of each tool. “×” means that TIMER was not suitable for estimating the cell fraction of the two microarray datasets (PBMC: GSE65133 and FL: GSE65136).

Meanwhile, we used two microarray datasets (GSE65135^[12]^ and GSE65133^[12]^ from GEO), which are obtained from disaggregated lymph node biopsies from patients with follicular lymphoma and samples from peripheral blood with immune cell ratios determined by flow cytometry. The abundance of each cell type measured by ImmuCellAI showed overall high positive correlations with the flow cytometry results in both datasets (Figure 2D and Figure S2B). In addition, ImmuCellAI showed the least correlation deviation in both RNA-Seq and microarray datasets (Figure 2E). The performance evaluation results indicated that ImmuCellAI has the best performance in both microarray and RNA-Seq data with stable and high precision in terms of estimation of abundance of the 24 immune cell types.

### 2.3. Case study of ImmuCellAI application for cancer immunotherapy response prediction

To investigate the impact of immune cell abundance on cancer immunotherapy, we applied ImmuCellAI to an anti-PD1 dataset, GSE91061^[21]^, which comprised 58 melanoma samples from a clinical trial on anti-PD1 therapy. We analyzed the results using two comparisons: responders vs. non-responders and on-treatment vs. pre-treatment. The abundance of three immune cells including Tc, γδ T and DC cells significantly increased with anti-PD1 treatment (Figure 3A; Mann–Whitney *U*-test, *p* < 0.05). Besides, Tc, γδ T and DC cells also significantly infiltrated more in responders compared with non-responders at the on-treatment time point (Figure 3B; Mann–Whitney *U*-test, *p* < 0.05). The results suggested that ImmuCellAI can provide important insights on the dynamic immune cell infiltration during immunotherapy and offer valuable indicator for immunotherapy response during the treatment.

**Figure3.**
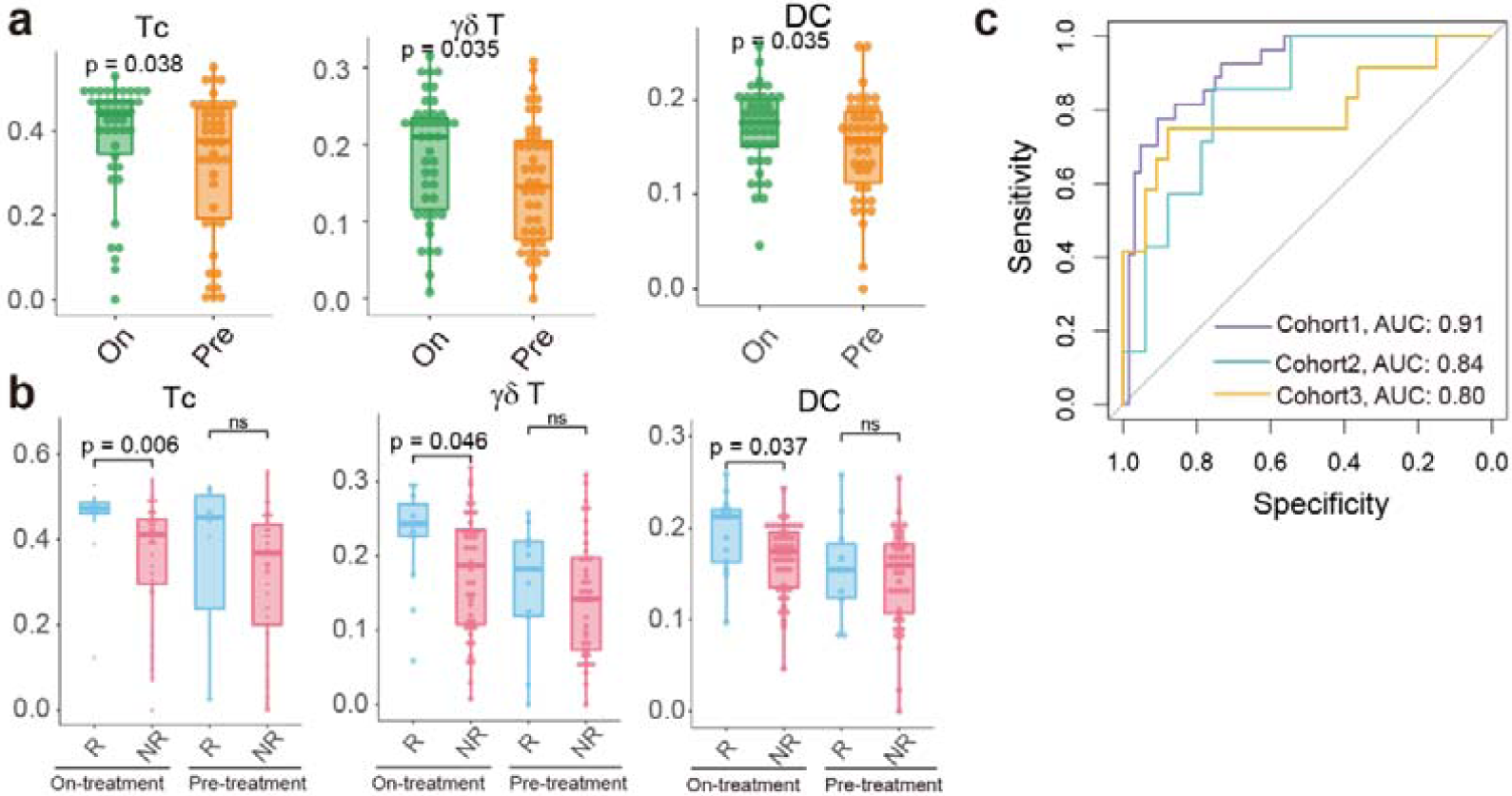
Case study of the application of ImmuCellAI to the immunotherapy datasets. A-B. The significant abundance differences of 3 types of immune cells in before (Pre) and during (On) anti-PD1 treatment (A), and responders (R) and non-responders (NR) at on-treatment (anti-PD1) time point. C. The receiver operating characteristic (ROC) curve of the immunotherapy response prediction model in the test and validation cohorts. “Cohort 1” contains 53 samples (8 responders and 45 non-responders, random sampling from GSE91061, GSE78220 and GSE115821). “Cohort 2” contains 41 samples (7 responders and 33 non-responders, SRP011540). “Cohort 3” contains 45 samples (12 responders and 33 non-responders, ERP107734).

Next, we applied the abundance of immune cells estimated by ImmuCellAI to predict the response to immune checkpoint blockade therapy. Five anti-PD1 or anti-CTLA4 therapy datasets (GSE91061^[21]^, GSE78220^[22]^, and GSE115821^[23]^, ERP107734^[24]^ and SRP011540^[25]^), involving a total of 176 patients, were analyzed, in which the former three datasets from GEO were used for training and testing in a support vector machine model based on the abundance of immune cells, and the last two cohorts from dbGAP were used to further validation the model (detail in methods). A feature integrating the abundance of 22 immune cell types (B cell, CD4^+^ naïve, CD8^+^ naïve, Tcm, Tc, DC, γδ T, Tem, Tex, iTreg, macrophage, MAIT, monocyte, neutrophil, NK, NKT, nTreg, Tfh, Th1, Th17, Th2, and Tr1) had the best performance in predicting immunotherapy response in the test data [area under curve (AUC) = 0.91] and other two validation cohorts (AUC 0.84 and 0.80) (Figure 3C). Overall, the abundance of immune cells measured by ImmuCellAI was highly predictive of immune therapy sensitivity (Figure 3C), suggesting that ImmuCellAI can serve as an ideal method for immunotherapy studies. We implemented the model for immune therapy response prediction as a functional module on the ImmuCellAI server.

### 2.4. Case study of ImmuCellAI application to TCGA pan-cancer data for predicting the infiltration of immune cells and patient survival

Increasing evidence has demonstrated that immune cells are critical in cancer progression, and the infiltration of different T-cell subsets could dramatically influence the treatment strategy and prognosis^[26]^. In this study, to demonstrate the application of ImmuCellAI to cancer data, we analyzed 17 cancer types in TCGA with gene expression data of both the tumor and adjacent tissues to survey the infiltration of immune cells. Partial correlation analysis was implemented to reduce false correlation, which may be caused by other factors, such as age and gender (Figure S3). The results indicated that the abundance of many immune cell types was significantly different (FDR < 0.1) between the tumor and adjacent tissue samples in most cancers, particularly for Tc, NK, NKT, Th2, iTreg, nTreg, and DC (Figure 4A). The iTreg, nTreg, Tr1, and monocyte cells were markedly enriched in the nidus of most cancer types, which is consistent with their immunosuppressive properties (Figure 4A). In contrast, several antitumor cells, such as γδ T, MAIT, NK, NKT, and Th2 cells, showed higher infiltration in adjacent tissues of most cancers, indicating that the tumor microenvironment may prohibit their access to the nidus.

**Figure 4.**
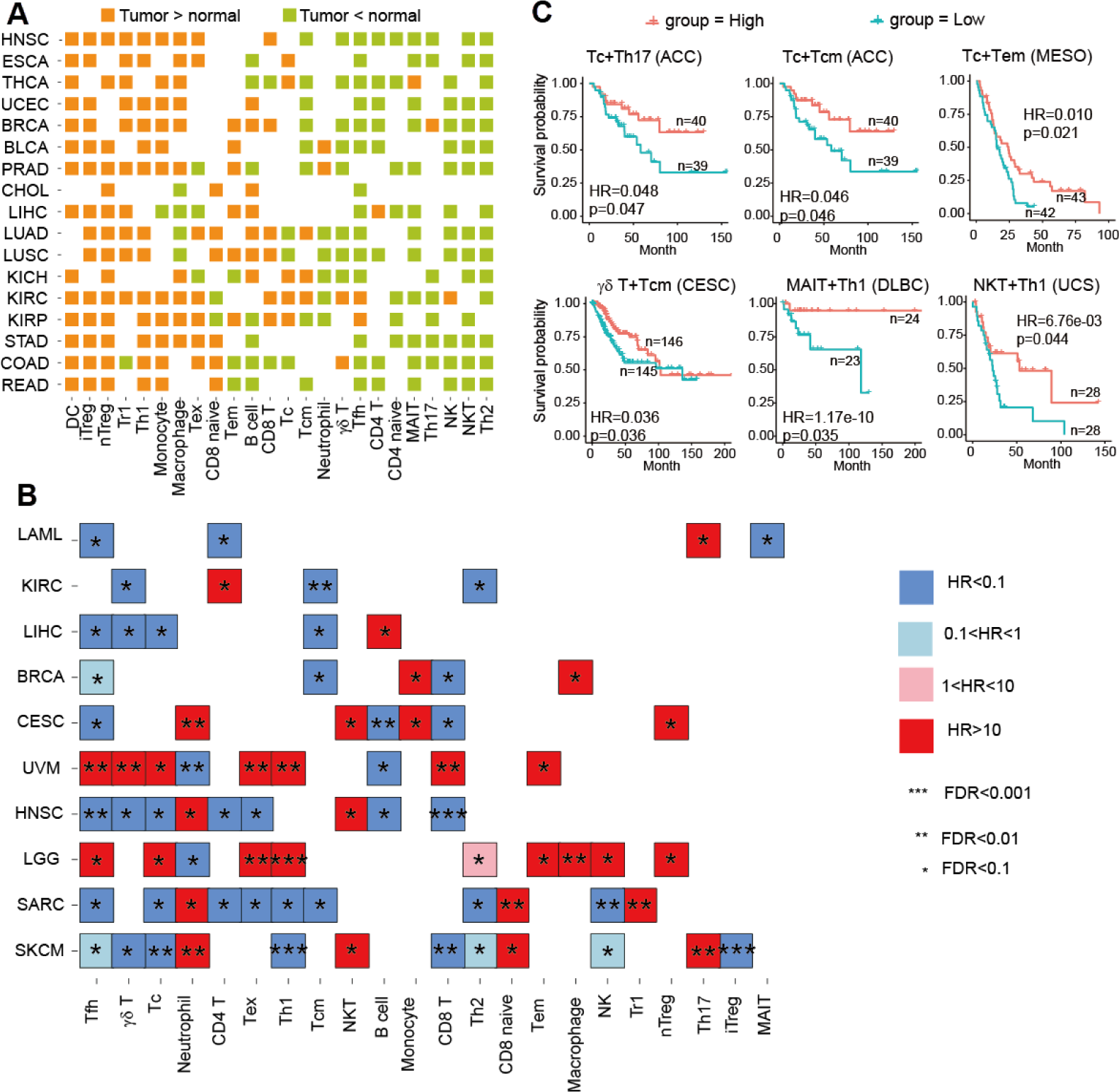
Analysis of the infiltration of immune cells in TCGA data by ImmuCellAI. **A.** A landscape of the comparison of the infiltration of immune cells between the tumor and adjacent tissues. The orange blocks indicate that cells infiltrated more in the nidus tissue and green blocks indicate the opposite. Statistical significance was evaluated using Wilcoxon’s rank sum test with an FDR of 0.10. B. Association of tumor-infiltrating immune cells with patient survival. For each cancer type, multivariate Cox regression was performed, with covariates including the abundance of immune cell, patient age at diagnosis, gender, and clinical stage. C. Kaplan–Meier curves of cancers by the combination of multiple immune cell types. Statistical significance and hazard ratios were calculated using multivariate Cox regression.

Furthermore, we investigated the effects of the infiltration of immune cells on patient survival by controlling other factors (i.e., age, gender, and stage). In a nutshell, the infiltration of most immune cell types significantly affected the overall survival of patients in different cancers (Figure 4B). The infiltration of most immune cells had opposite effects on survival in LGG and UVM compared with that in other cancers (Figure 4B). Furthermore, skin cutaneous melanoma (SKCM) had the most immune cell types (12/24) significantly associated with patient survival (FDR <0.1; Figure 4B and Figure S4). The infiltration of T-cell subsets (e.g., γδ T cells, Th1, Th2, and iTreg) had positive effects on long-term survival in patients with SKCM, whereas patients with high infiltration of CD8^+^ naïve and neutrophils were associated with worse outcomes (Figure 4B and Figure S4). Although the infiltration of a single immune cell type in some cancer types was not related with patient survival, the infiltration of a combination of multiple immune cell types was significantly associated with survival (multivariate Cox regression, *p* < 0.05), such as Tc + Th17 in ACC, Tc + Tcm in ACC, Tc + Tem in MESO, γδ T + Tcm in CESC, MAIT + Th1 in DLBC, and NKT + Th1 in UCS (Figure 4C). In addition, the infiltration of immune cells was correlated with microsatellite instability (MSI) after partial correlation analysis was performed to reduce false correlation caused by other features, such as age and gender (Figure S5A and S5B). In colon adenocarcinoma (COAD) and stomach adenocarcinoma (STAD), patients with high-MSI cancer (MSI-H) showed a significantly higher infiltration of antitumor and tumor helper cells, such as Tc, γδ T, NK, and DC (Figure S6A), but a significantly lower infiltration of tumor suppressor cells (Tr1 and neutrophils) and CD8 naïve cells (Figure S6b). These factors may partially explain the better outcomes of patients with MSI-H colorectal cancer undergoing immunotherapy^[27]^. Furthermore, we observed that some immune cell types showed stage-related profiles in cancers. For example, the infiltration of Tex, Th1, iTreg, and CD8^+^ T cells gradually increased with the development of KIRC (Figure S7).

## 3. Discussions

Increasing evidence suggests that immune cells play critical roles in carcinogenesis and progression, and a proper proportion of T-cell subsets could contribute to long-term clinical benefits of anticancer treatments^[28]^. Investigating the abundance of immune cell types could provide insights into a more comprehensive understanding of the immune status of patients and could thus benefit disease therapy^[29]^. In this study, we developed ImmuCellAI, a highly accurate method of estimating the abundance of immune cells, particularly T-cell subsets, from transcriptome data. The case study application results on immunotherapy and pan-cancer data suggest that ImmuCellAI is a very useful tool in cancer immunology.

ImmuCellAI uses a global enrichment algorithm to enumerate immune cells based on transcriptome data and shows a robust and accurate performance. Although there are several other methods available for immune cell identification, most of them, with the exception of xCell, can only roughly identify very few T cell subsets. The unique function of ImmuCellAI is that it can accurately estimate the abundance of different T-cell subsets, which is particularly important in cancer therapy. For those immune cells that could be identified by other methods, our comparison results showed that ImmuCellAI had the highest consistency with flow cytometry results for most cells (Figure 2B, 2C, 2D and 2E). Because there are very limited T-cell subsets with both flow cytometry data and RNA-Seq data, we produced two datasets using flow cytometry analysis for all 24 immune cell types and sequenced their RNA. The results confirmed that ImmuCellAI has the best performance in terms of accurately identifying these T-cell subsets, which is an advantage of this method.

Tumor-infiltrating T cells could serve as a prognostic factor and predictor of therapeutic efficacy^[30]^. In our result, the abundance of Tc, γδ T and DC cells were significantly increased in both comparisons of on-treatment vs. pre-treatment and responders vs. non-responders (Figure 3A and 3B). DC cells are antigen-presenting cells (APC) that are essential for the activation of immune responses, which have the potential to turn immunologically “cold” tumors into “hot” tumors^[31]^. Tc cells play key roles in the tumor cell killing process^[32]^, and γδ T cells also function in the antigen recognition and tumor killing process^[33]^. To extend these results, an immunotherapy model was proposed, taking the immune cell abundance of pre-treatment samples into account. The model achieved a high accuracy for immunotherapy response prediction with an average AUC of 0.85 (Figure 3C), suggesting that it is a promising tool in immunotherapy studies; this model was implemented for immunotherapy response prediction on the ImmuCellAI server. This is the second user-friendly web server for immunotherapy response prediction, except for the TIDE^[34]^. Comparing with the reported accuracy of TIDE, our ImmuCellAI has a little bit higher accuracy. In addition, to date, most studies have focused on the infiltration of CD8^+^ T cells as a predictive biomarker for response to immune checkpoint blockade therapy^[35]^. Our study indicated that integrating many immune cells, particularly different T-cell subsets, could serve as a biomarker for better therapy response prediction (Figure 3C). Thus, the systematic evaluation of immune cell abundance could be an effective approach for predicting immune checkpoint blockade therapy response and improving the effects of cancer immunotherapy^[36]^. Some of the results of the infiltration of immune cells in TCGA cancer data were consistent with those of previous reports, for example, the infiltration of Tc cells is more often observed in kidney cancer but less so in colorectal cancer^[37]^. These results indicate the powerful and unique function of ImmuCellAI on cancer immunology and immunotherapy research.

Although ImmuCellAI had the best performance in comparison with other methods, it still has several limitations that need to be addressed. First, ImmuCellAI could only estimate the relative abundance of immune cells based on the deviation of gene signatures. It could not provide the absolute amount of each immune cell type. Moreover, ImmuCellAI did not consider the spatiotemporal localization of immune cells and the abundance of cancer cells. Furthermore, ImmuCellAI lacked the sensitivity to identify cell types with low abundance (e.g., NK cells in samples from EPIC). In addition, the sample size used in the immunotherapy case study was relatively small, and the performance of our model needs to be tested in larger cohorts. Other immune cell subsets, besides the T cells used in our method, also need to be tested in future studies.

In summary, this study presented an accurate and reliable tool ImmuCellAI to dissect T-cell properties and explore the infiltration of immune cells in cancer. The best advantage of ImmuCellAI is its ability to accurately estimate the abundance of 18 T-cell subsets, which is its unique function. The results of ImmuCellAI provided valuable prognostic predictors and comprehensive resources to elucidate cancer–immune interactions, which could facilitate applications of cancer immunotherapy and precision medicine.

## 4. Materials and Methods

### 4.1. The main algorithm of Immune Cell Abundance Identifier (ImmuCellAI)

The main algorithm of ImmuCellAI, presented in Figure 1B, includes three main steps: (1) reference expression matrix (RT) and marker gene preparation, (2) enrichment score calculation, and (3) compensation matrix correction.

#### 4.1.1. Reference expression matrix and marker gene preparation

The datasets of the expression profiles of 24 immune cell types (Figure 1A) were downloaded from the National Center for Biotechnology Information Gene Expression Omnibus (GEO) database. In total, 415 datasets from 26 studies were manually curated to build RT of the immune cell types (table S2). Gene expression data was obtained from CEL files according to the frozen robust multiarray analysis protocol with batch effect correction^[38]^. Each line of the matrix denotes the expression of a gene in the 24 immune cell types. The median value was used if there were multiple samples of a cell type.

Furthermore, we developed a gene signature for each cell type by integrating the marker genes obtained from the literature and other analytical methods, such as CIBERSORT and xCell; thus, a total of 2547 genes were collected (denoted as Ga, table S1). Next, a robust marker gene set per immune cell type was selected using *in silico* simulated data taking advantage of the TCGA data, which was based on the work of Li et al.^[14]^ For each cancer type in the TCGA data, the expression of Ga in the samples (log2 transferred) and immune cell reference profiles were used to simulate the immune cell infiltrated tumor samples with known fractions. To control the mixing ratios of immune cell components for maintaining the correlative structure of real data, we first calculated the gene–gene covariance matrix Σa for all genes in Ga using tumor expression data. Then, we randomly sampled 24 numbers (f1–f24) from Uniform (0,1) and calculated μa (length *n*), which is the average of gene expression in the reference profiles of the 24 immune cell types weighted by f1–f24. Next, we sampled a vector of length *n* from the multivariate normal distribution with mean μa and covariance Σa. For each cancer type, we simulated the same number of samples as its sample size in the TCGA data.

Then, for all collected marker genes in Ga, we calculated the average correlation between gene expression in simulated samples with cell fractions using Pearson correlation for all cancers, and genes with an average correlation of *r* ≥ 0.6 were selected (denoted as G1). Next, for each marker gene per immune cell, the standard correlation deviation among the cell with other cells was calculated, and genes with a standard deviation of >1.5 were selected (denoted as G2). The deviation between CD4^+^ T and CD4^+^ T-cell subsets (such as CD4^+^ naïve and Th1) as well as that between CD8^+^ T and CD8^+^ T-cell subsets was not calculated. Finally, a robust marker gene set per immune cell type was obtained by intersecting G1 with G2 (denoted as Gf), which included 344 marker genes of the 24 immune cell types (table S1). In addition, we constructed a sparse matrix (ST) for these marker genes in which “1” means that the gene is a marker gene in the corresponding cell type.

#### 4.1.2. Enrichment score calculation

For a user-uploaded expression dataset, ImmuCellAI first calculates the expression deviation of all marker genes compared with RT. Here two different approaches were implemented to deal with the microarray and RNA-Seq datasets.

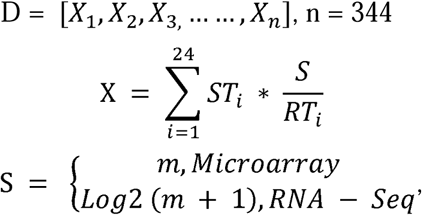

where vector D denotes the relative deviation of marker genes and ST_*i*_ is the vector in ST for marker genes of cell type *i. RT*_*i*_ is the reference marker gene expression in cell type *i*, whereas *S* indicates the gene expression in the user-provided dataset.

The single sample gene set enrichment analysis (ssGSEA) algorithm in the GSVA package^[39]^was used to estimate the abundance of immune cell types. The ssGSEA enrichment score for deviation vector D of the gene signature of each immune cell type (named ES) was used to indicate the relative abundance of immune cell types in the user-provided dataset. A higher enrichment score indicates a higher abundance of the immune cell type in the mixture sample than that of other cell types.

#### 4.1.3. Compensation matrix correction

Some immune cell types may share a part of common marker genes, which will cause bias in the estimation of abundance of these immune cell types. Thus, ImmuCellAI used a compensation matrix and least square regression method based on the work of Aran et al.^[11]^ to fix this issue. After the estimation of abundance of the detected immune cell types in a dataset, we reassigned the weights of common marker genes for these immune cell types with the following steps: (1) A N * N contribution matrix was produced by calculating the mutual contributions of marker genes in RT using ssGSEA. (2) Each column of the contribution matrix was divided by a diagonal value and weighted by the proportion of non-diagonal elements, and a compensation matrix was obtained (named C). (3) To reduce redundancy and overestimation of compensation between detected immune cell types, ImmuCellAI discarded the compensational calibration between the parental immune cell type and its subsets (e.g., CD4^+^ T and Th1 cells), and limited the total compensation level at 0.5 for the non-diagonal cell types. (4) The least square method was used to calibrate the enrichment score based on the compensation matrix C.

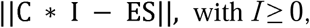

where the parameter ES is the ssGSEA enrichment score of detected immune cell types and C is the compensation matrix. Finally, after calibration, we deemed the abundance of 24 immune cell types (named I) to be high confidence.

### 4.2. Benchmark dataset preparation

#### 4.2.1. Our datasets for the 24 immune cell types

Heparinized blood samples from seven patients with leukemia and five healthy adult volunteers were collected from Wuhan Central Hospital, China. Fresh blood samples were treated with Pharm Lyse (BD Biosciences, San Jose, CA, USA) to remove erythrocytes. Cells from each sample were used in parallel experiments of flow cytometry and RNA extraction. This study was approved by the ethics committee of Tongji Medical College, Huazhong University of Science and Technology, and followed the Declaration of Helsinki principles. The proportions of the 24 immune cell types used in our study were examined by flow cytometry using the combined markers listed in table S3 and antibodies listed in table S4. All antibodies were purchased from BD Biosciences, except those used against TCR-Vβ2 and TCR-Va7.2 (Miltenyi Biotec, Bergisch Gladbach, Germany).

The total RNA extracted from the cells of all 12 samples was used for RNA sequencing (RNA-Seq) (PE150) via the Illumina HiSeqTM4000 platform by Haplox (Jiangxi, China). RNA-Seq reads were mapped to Ensembl v81 (GRCH38) and processed using the HISAT2-StringTie-ballgown pipeline. We used fragments per kilobase per million mapped reads to calculate gene expression levels.

#### 4.2.2. Other public datasets

The microarray datasets and corresponding flow cytometry results were obtained from GEO (accession nos. GSE65135 and GSE65133), which include 14 disaggregated lymph node biopsies from patients with follicular lymphoma and 20 peripheral blood samples from individuals vaccinated for influenza, respectively.

The RNA-Seq dataset from EPIC contains both RNA-Seq data and their corresponding flow cytometry results of five immune cell types (B, CD4^+^ T, CD8^+^ T, NK, and cancer cells) from four patients with melanoma^[13]^. Because of the scarcity of bulk RNA-Seq datasets containing both gene expression profiles and flow cytometry counts for different immune cell types, particularly for T-cell subsets, we simulated two bulk RNA-Seq datasets by integrating the expression profiles of seven cell types (CD4^+^ naïve, CD8^+^ naïve, MAIT, Tcm, Tex, Treg, Th) from single-cell RNA sequencing data. The transcripts per million (TPM) normalized expression of liver and lung cancers from two *Nature* papers were collected from GEO (GSE98638^[18]^ and GSE99254^[19]^). Based on the work of Max et al.^[40]^, single-cell expression was normalized as follows: for each single-cell dataset, the TPM values were transformed to

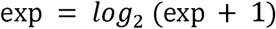

To ensure cross-sample comparability, the expression of all single-cell samples from the same dataset were normalized to the average expression of 3686 housekeeping genes^[41]^ as follows:

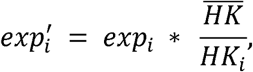

where *exp*_*i*_ represents the gene expression profile of sample *i, HK*_*i*_ denotes the average gene expression of all housekeeping genes in sample *i*, and 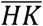 is the average expression of all housekeeping genes in all samples.

A single-cell sequencing dataset from 19 patients with melanoma was collected from GEO (accession GSE72056), which is the normalized expression matrix as described above by Tirosh et al.^[20]^ and contains the single-cell RNA-Seq of B cells, T cells, macrophages, NK cells, and three other nonimmune cell types. Because CD8^+^ and CD4^+^ T cells can be easily distinguished by CD4, CD8A, and CD8B expression, we divided T cells into CD8^+^ T cells, CD4^+^ T cells, and others. Then, the bulk expression of each sample was identified by aggregating normalized expression from all cell barcodes for each patient sample. The cell ratio per cell type in a sample was calculated by the cell number of a specific cell type divided by the total number of cells (table S5–S7).

### 4.3. Performance assessment of ImmuCellAI

The performance of ImmuCellAI was evaluated using both microarray and RNA-Seq datasets and compared with that of five other methods (CIBERSORT, EPIC, MCP-counter, TIMER, and xCell). For a given immune cell type, the accuracy and sensitivity of each method were measured using the Pearson correlation between the results of *in silico* method and flow cytometry counting in samples (named *r*_*i*_). In addition, we introduced the correlation deviation to measure the global performance of each method, which took the sample size and overall accuracy into consideration.

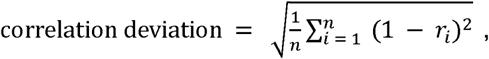

where *n* is the amount of immune cell types detected in samples and *r*_*i*_ is the Pearson correlation of immune cell type *i*.

### 4.4. Case study of immune therapy and prediction model building

Five immune checkpoint therapy datasets, including those from anti-PD1- or anti-CTLA4-treated patients with melanoma or gastric cancer, were collected from the GEO database (GSE91061^[21]^, GSE78220^[22]^, and GSE115821^[23]^) and dbGAP (ERP107734^[24]^ and SRP011540^[25]^). The abundance of infiltrating immune cells was calculated by ImmuCellAI and used to build the response prediction model.

The immunotherapy response prediction model was built using support vector machine with the radial basis function kernel. The training features were the abundance of immune cell types. The sequential backward feature selection algorithm was used to minimize the feature number and improve the performance. At first, three GEO datasets composed of 91 pre-treatment samples (response: complete response and partial response, n = 27, non-response: stable disease and progressive disease, n = 64) were used to train and test the model. The undersampling method was used to fit the unbalanced sample size between responders and non-responders with 38 samples in the training and validation cohort (19 responders and 19 non-responders, 5 fold cross validation) and 53 samples in the test cohort (8 responders and 45 non-responders). Then, the other two cohorts from dbGAP (ERP107734: 12 responders and 33 non-responders; SRP011540: 7 responders and 33 non-responders) were used to further validate the model. The area under curve (AUC) was used to measure the model performance.

The gene expression profiles of TCGA samples and the clinical information were downloaded from Broad GDAC Firehose (https://gdac.broadinstitute.org/).

### 4.5. Statistical analysis

Basic statistical analyses, such as Wilcoxon rank sum test and Pearson correlation, were performed using R language. The correlations between clinical indicators and the abundance of immune cell types were evaluated using partial correlation analysis in the R package “ppcor.” Multivariate Cox regression, log-rank test, and Kaplan–Meier in R package “survival” were used to assess the relationships between the abundance of immune cell types and survival time. The *p* values for each test were calibrated using FDR, and the FDR threshold was 0.1 in case studies. All results supported the current study and were deposited into the ImmuCellAI website (http://bioinfo.life.hust.edu.cn/web/ImmuCellAI/).

## Acknowledgments

We express our gratitude to all study participants.

## Authors’ contributions

YRM and QZ: Performed formal analysis, conceptualized, conceived method, and wrote the original draft and edited the manuscript; QL: Collected samples and performed the experiments; ML and GYX: Performed formal analysis; HXW: Provided sample and experiment assistance; AYG: Conceptualized, wrote-reviewed and edited the manuscript, and funded and supervised the study.

## Data and materials availability

The sequence data sets reported in this paper have been deposited in the National Genomics Data Center with accession no. CRA001839 and CRA001840.

## Table of contents

Immune Cell Abundance Identifier (ImmuCellAI) is a novel gene set signature-based method for precisely estimating the abundance of 24 immune cell types including 18 T-cell subsets. Application of ImmuCellAI to immunotherapy datasets revealed the dynamic change of immune cell abundance during immune checkpoint blockade therapy. An ImmuCellAI result-based model for predicting the immunotherapy response achieved high accuracy with AUC 0.80~0.91.

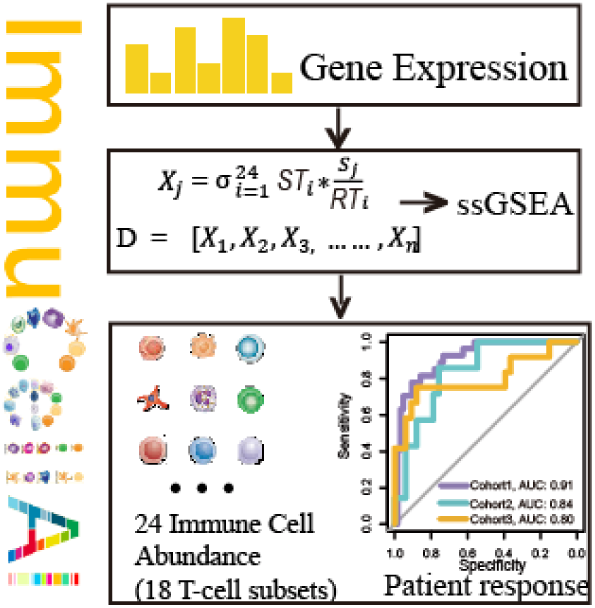

## Supporting Information

**Figure S1.**
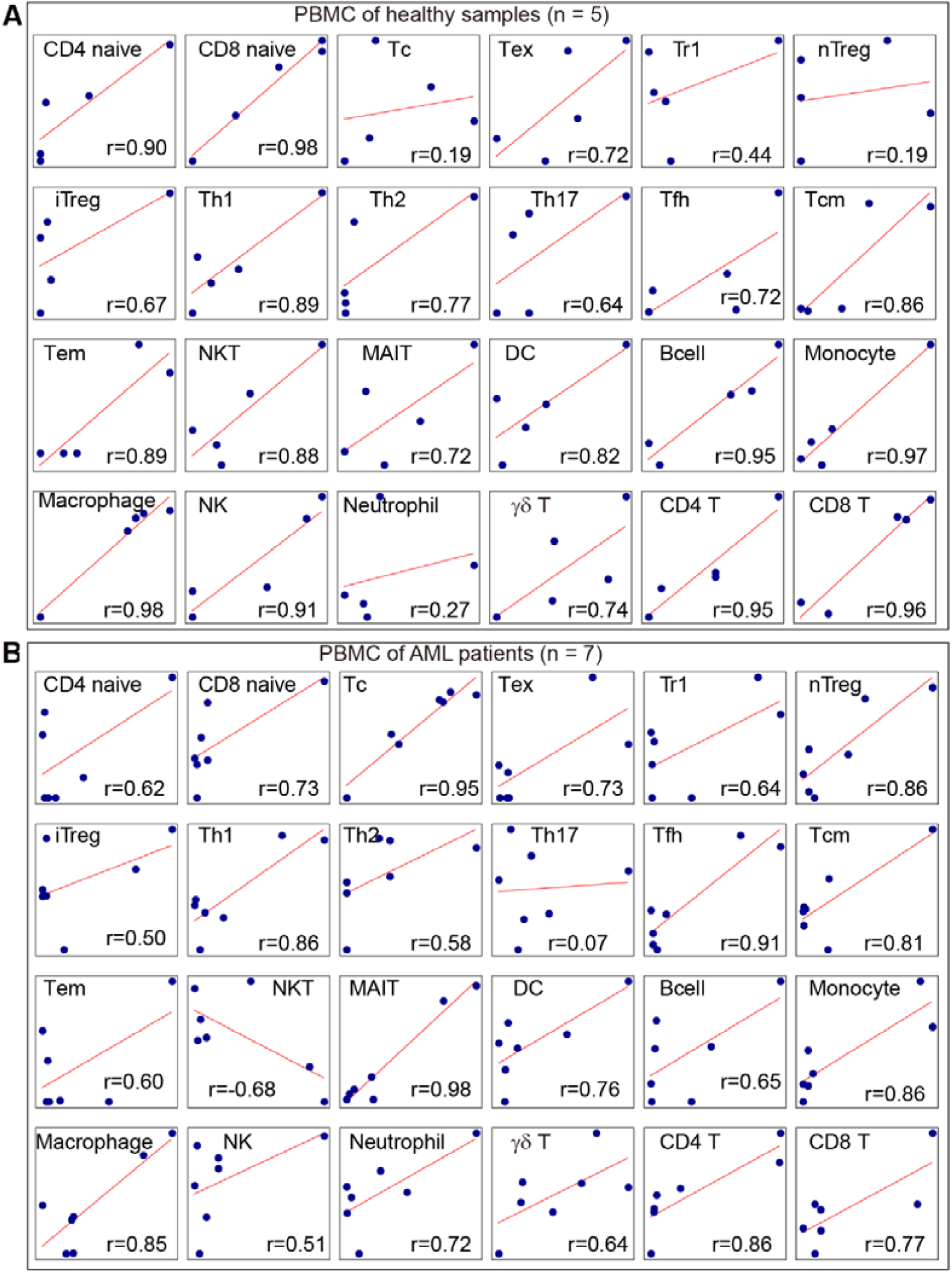
Direct comparison (Pearson correlation analysis) between cell fractions predicted by ImmuCellAI and the real cell abundance listed in Fig. 2B on PBMCs of samples from healthy individuals (A) and patients with AML (B).

**Figure S2.**
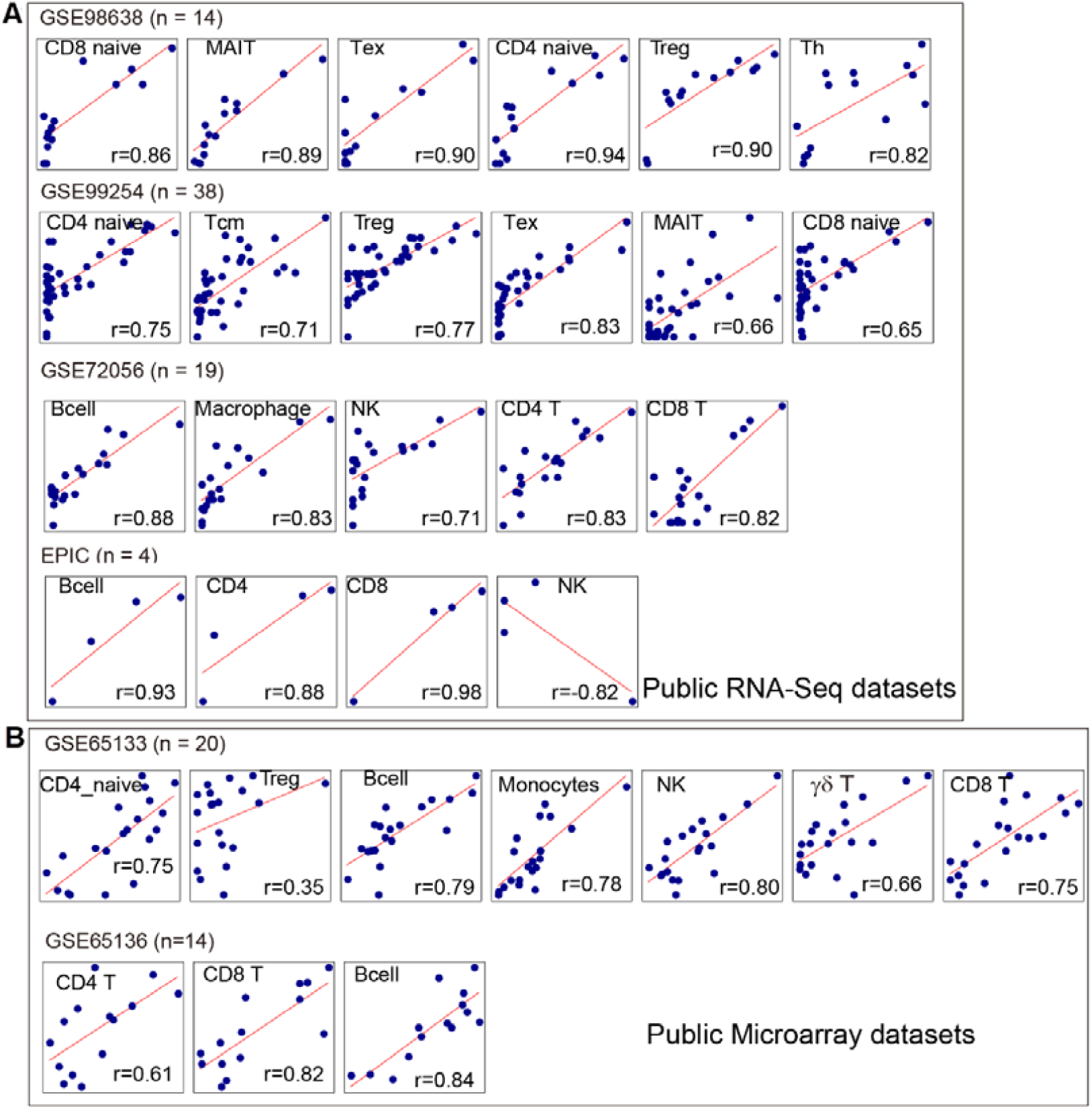
Direct comparison between cell fractions predicted by ImmuCellAI and the real cell abundance listed in Fig. 2C and 2D in public datasets estimated by RNA-Seq (A) and microarray (B).

**Figure S3.**
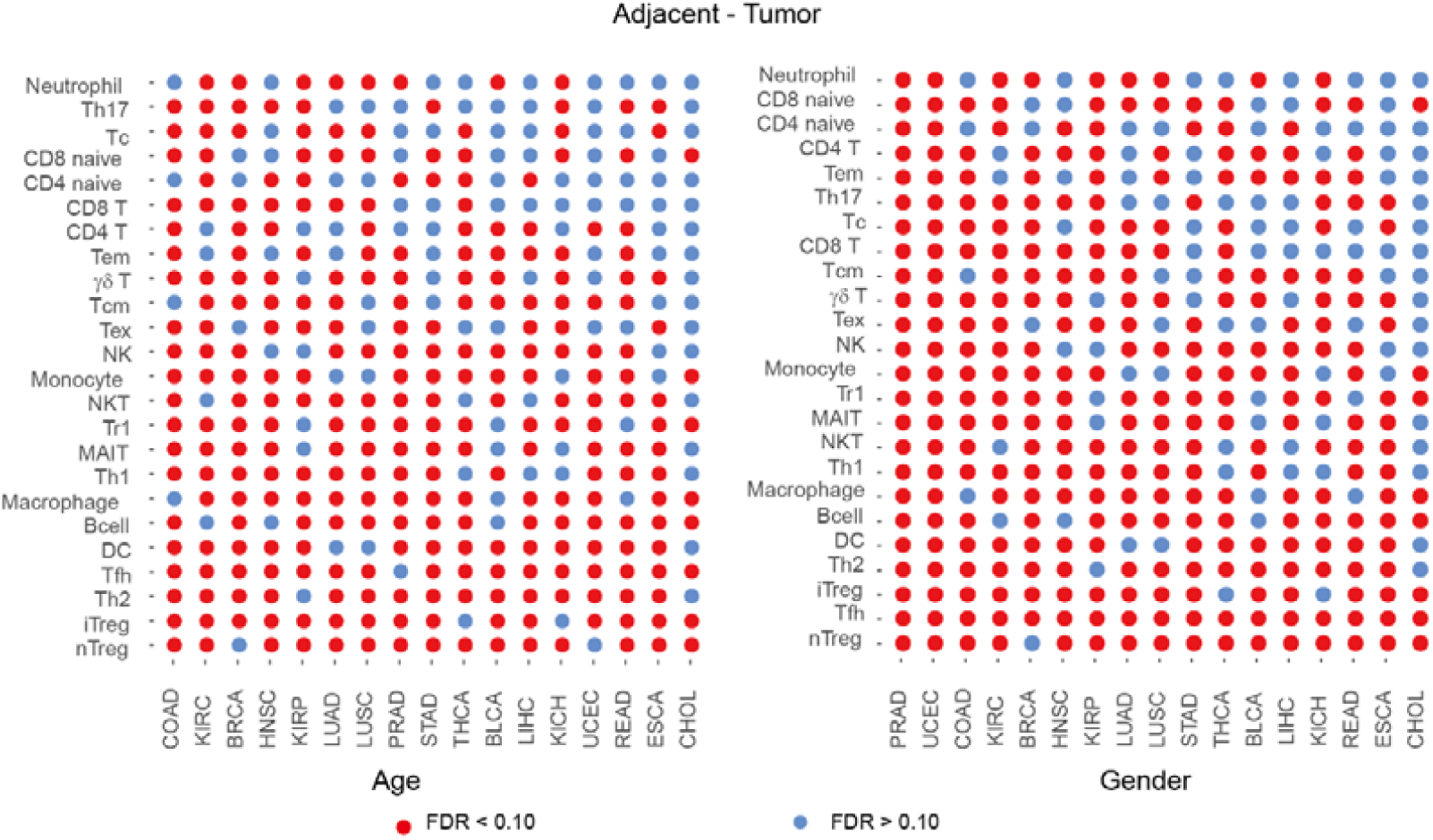
Partial correlation analysis of the infiltration of immune cells between the nidus and adjacent tissues, taking age and gender into consideration

**Figure S4.**
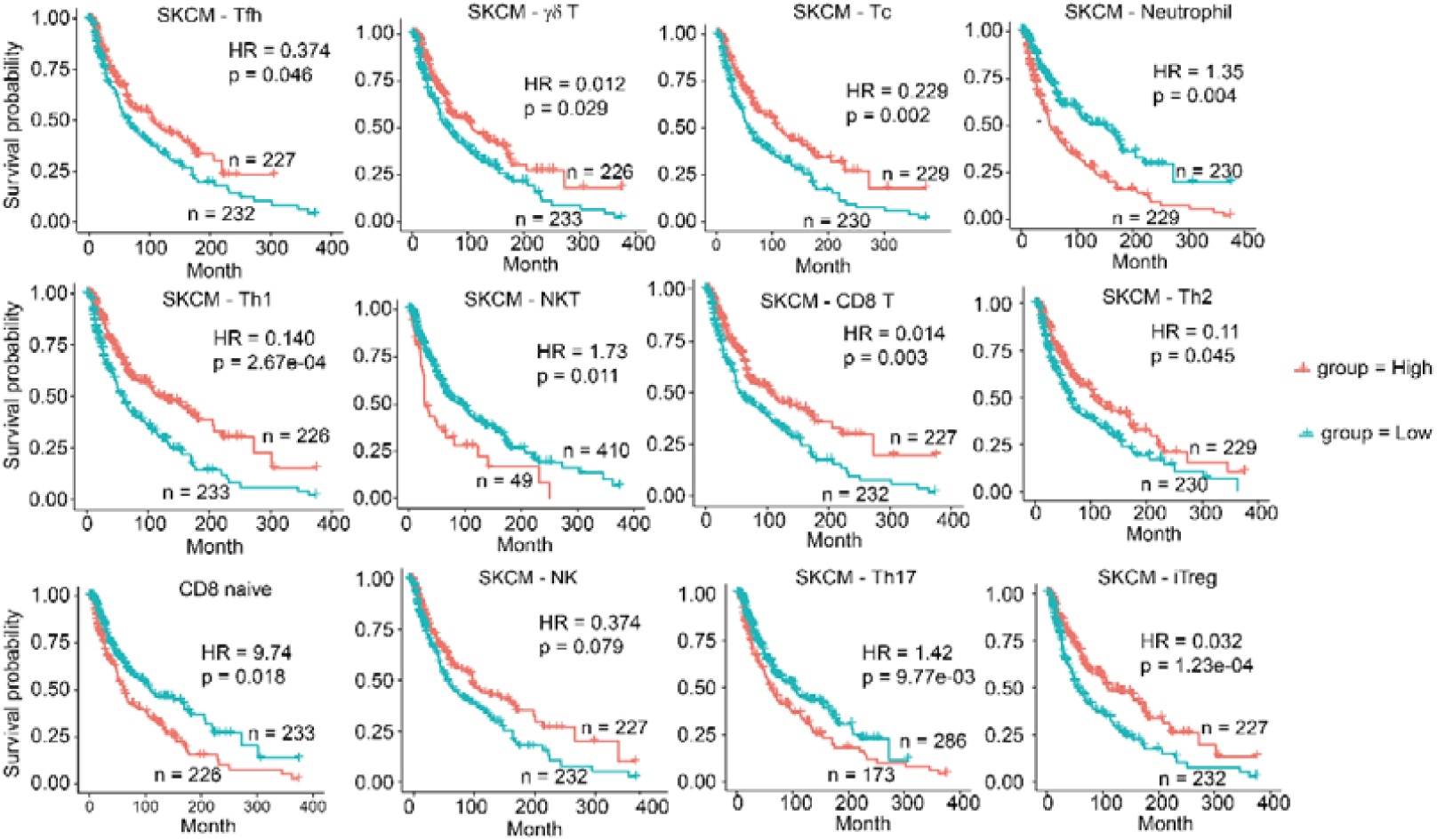
Kaplan–Meier curves of melanoma (SKCM) by the infiltration of 12 immune cell types including Tfh, γδ T, Tc, Neutrophil, Th1, NKT, CD8 T, Th2, CD8 naïve, NK, Th17, and iTreg.

**Figure S5.**
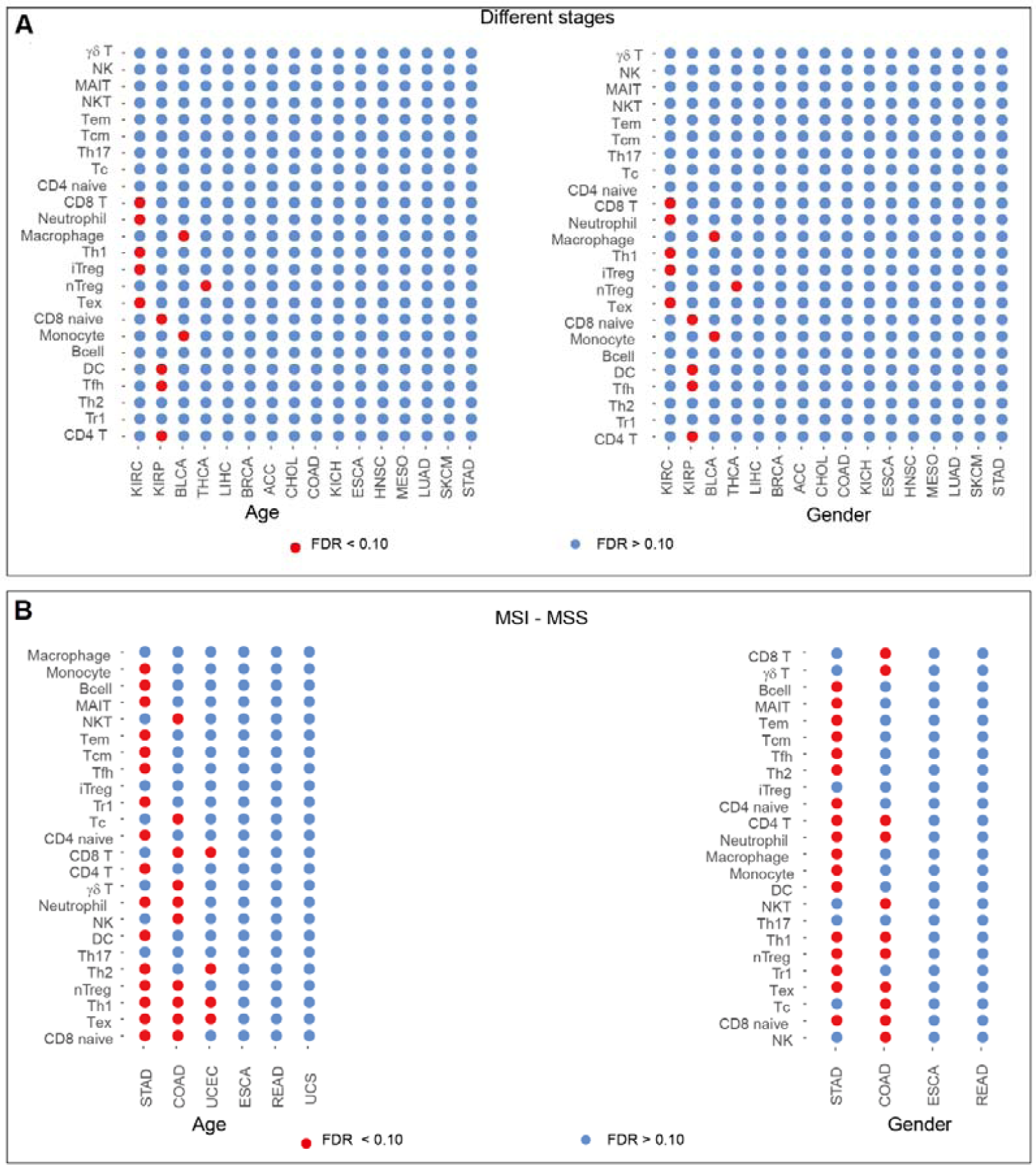
Partial correlation analysis of the infiltration of immune cells and clinical factors (age and gender)

**Figure S6.**
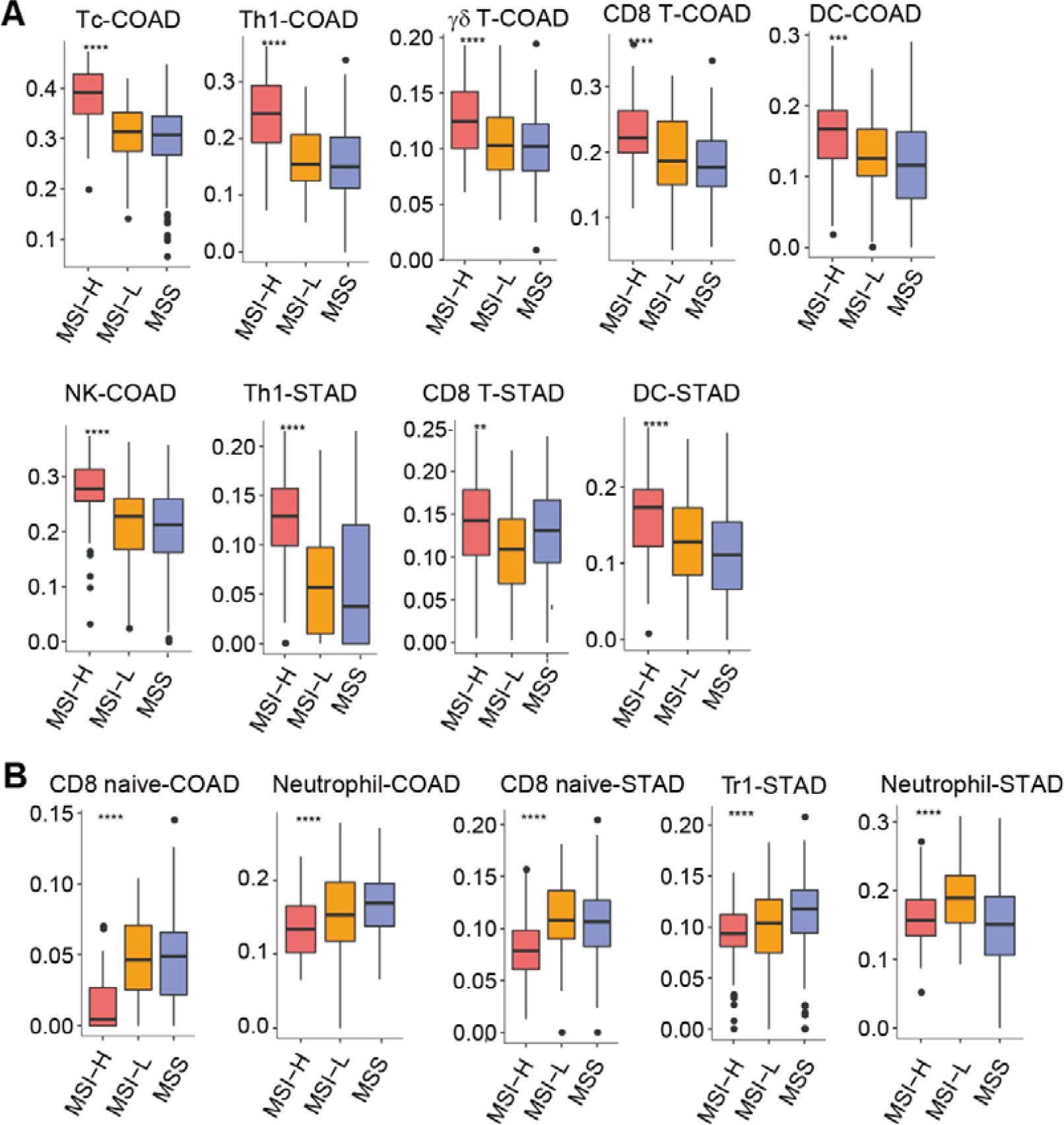
Association of the infiltration of antitumor (A) and tumor suppressor (B) immune cells with microsatellite instability (MSI) status in COAD and STAD.

**Figure S7.**
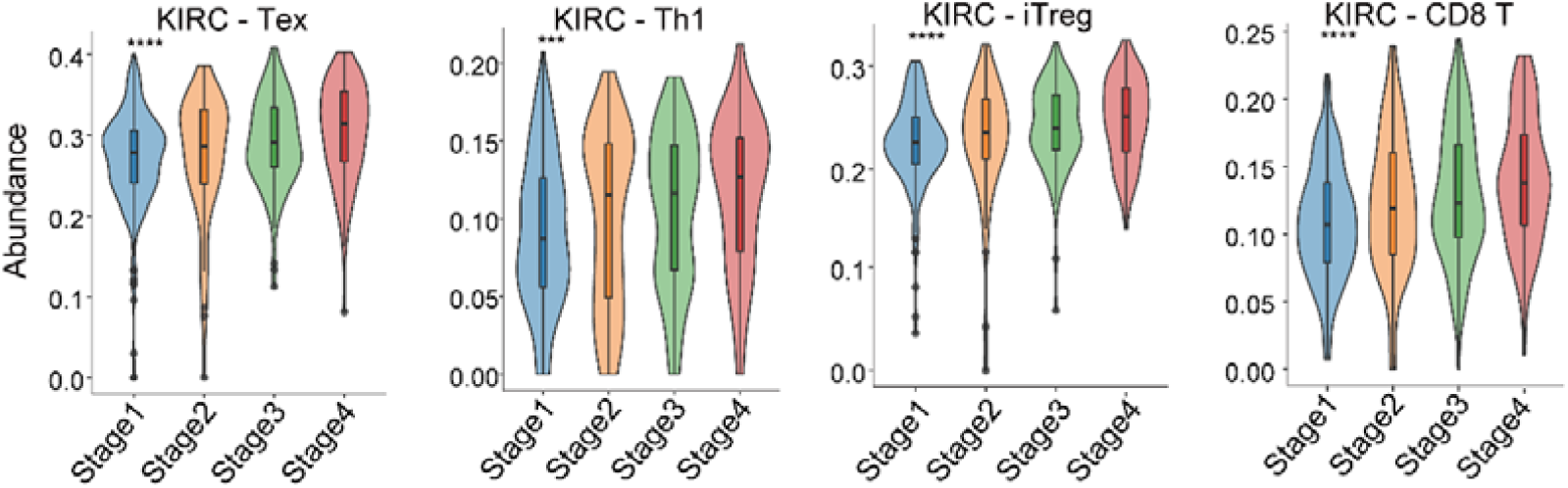
The abundance of tumor-infiltrating Tex, Th1, iTreg, and CD8^+^ T cells elevated with the clinical stage in KIRC

**Table S1** All marker genes used in ImmuCellAI

**Table S2** A list of immune cell reference expression profiles collected from the GEO database

**Table S3.** Immune cell antibodies used in flow cytometry.

| Cell type | Antibody |
| --- | --- |
| Bcell | CD45+CD19+ |
| DC | CD11c+HLA-DR+ |
| Macrophage | CD45+CD11b+ |
| Monocyte | CD45+CD14+ |
| Neutrophil | CD66b+CD11c+ |
| NK | CD45+CD56+CD3- |
| NKT | CD45+CD3+CD56+ |
| CD4 naïve | CD3+CD4+CD45RA+CCR7+ |
| CD4 T | CD3+CD4+ |
| iTreg | CD4+CD25+FOXP3+Helios- |
| nTreg | CD4+CD25+FOXP3+ Helios+ |
| Tfh | CD4+CD25-CXCR5+PD-1+ |
| Th1 | CD45+CD4+IFN-γ |
| Th17 | CD45+CD4+IL-17a |
| Th2 | CD45+CD4+IL-4 |
| Tr1 | CD4+CD25-LAG3+ |
| CD8 naïve | CD3+ CD8+CD45RA+ CCR7+ |
| CD8 T | CD3+CD8+ |
| Exhausted | CD45+CD3+CD8+PD-1+ |
| Cytotoxic | CD3+CD8+CD57+TCRVβ+ |
| Central memory | CD45+CD3+CD45RO+CCR7+ |
| Effector memory | CD45+CD3+CD45RO+CCR7- |
| Gamma delta T ( $\gamma\delta$T ) | CD3+TCRVδ2+ |
| MAIT | CD3+ CD161++ Va7.2 TCR+ |

**Table S4.**
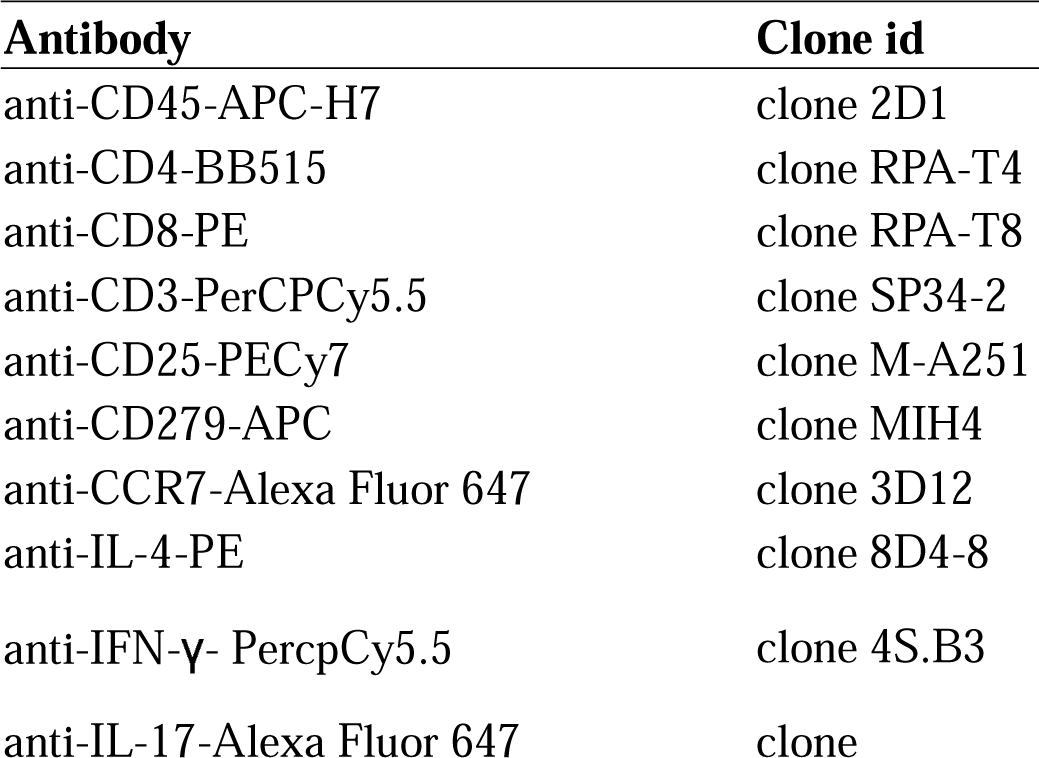
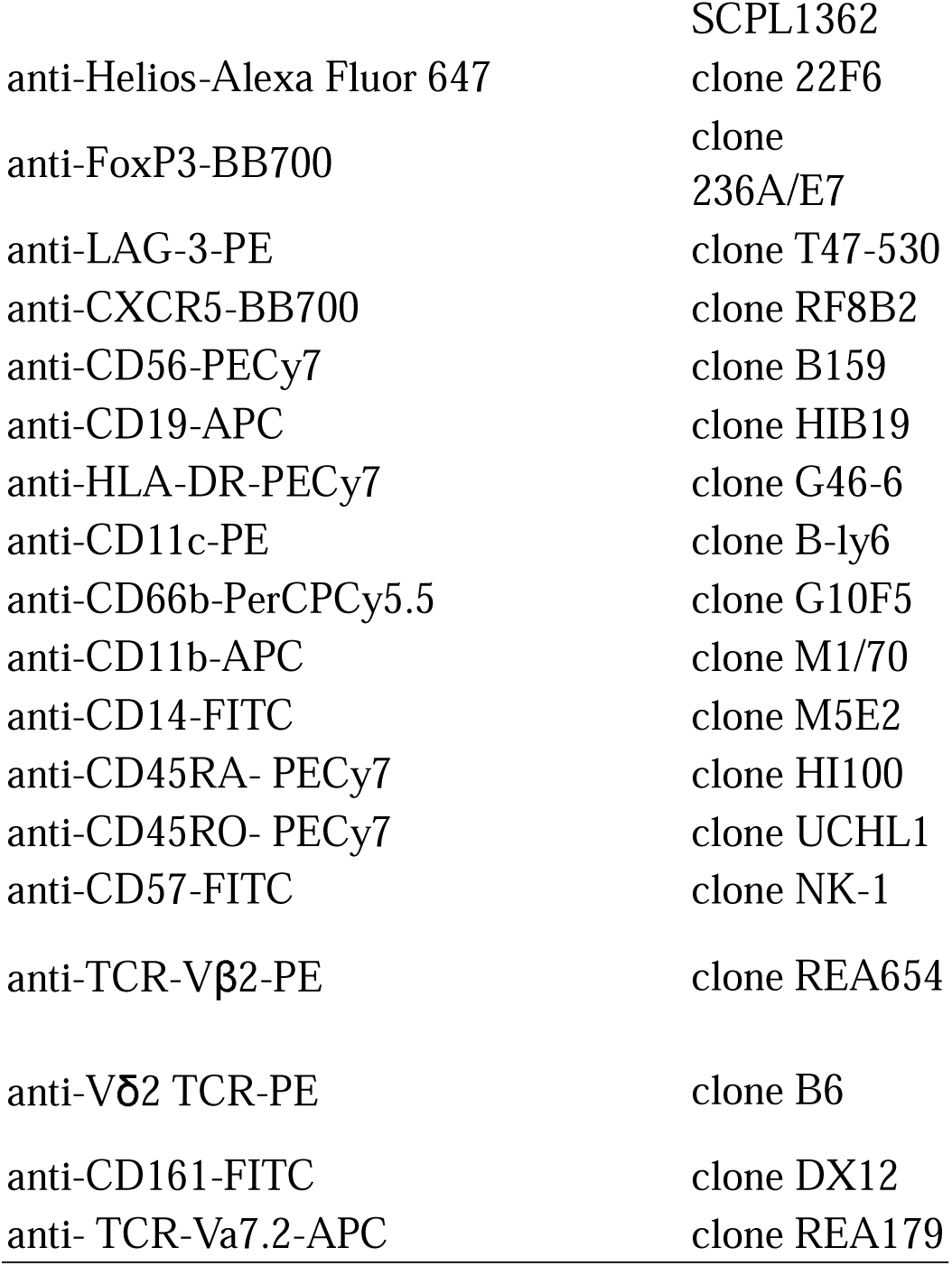
Antibody and its corresponding clone id used in flow cytometry.

**Table S5.**
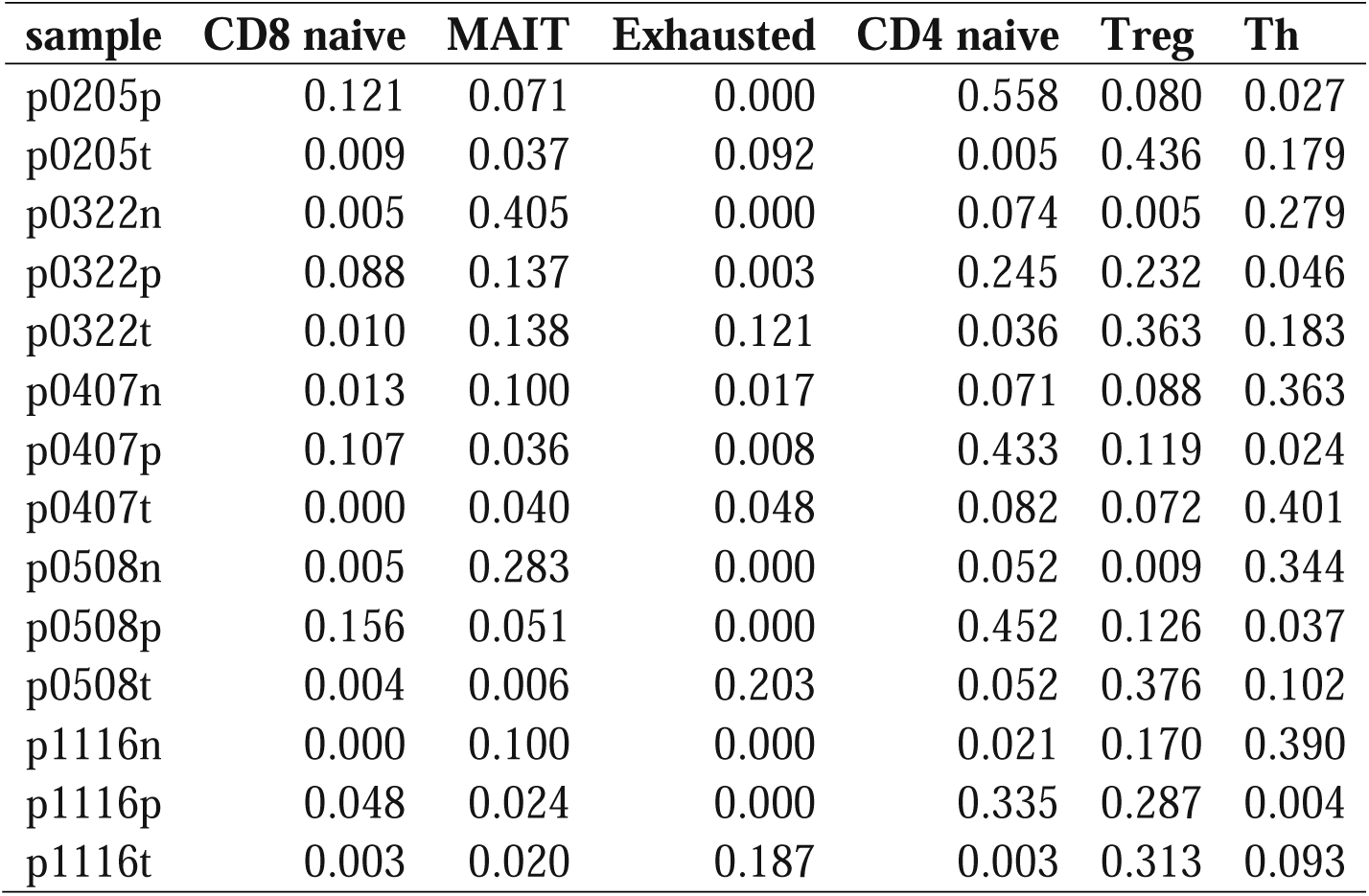
The real immune cell fraction calculated from single-cell barcode of GSE98638.

**Table S6.** The real immune cell fraction calculated from single-cell barcode of GSE99254.

| **Sample** | **CD4 naive** | **Tcm** | **Treg** | **Exhausted** | **MAIT** | **CD8 naive** |
| --- | --- | --- | --- | --- | --- | --- |
| P0616A_P | 0.158 | 0.055 | 0.060 | 0.000 | 0.004 | 0.070 |
| P0616A_T | 0.004 | 0.015 | 0.126 | 0.004 | 0.024 | 0.007 |
| P0616P_P | 0.154 | 0.101 | 0.000 | 0.000 | 0.016 | 0.202 |
| P0616P_T | 0.000 | 0.000 | 0.017 | 0.111 | 0.000 | 0.017 |
| P0617_N | 0.003 | 0.091 | 0.023 | 0.000 | 0.000 | 0.003 |
| P0617_P | 0.063 | 0.194 | 0.063 | 0.004 | 0.004 | 0.026 |
| P0617_T | 0.000 | 0.015 | 0.031 | 0.037 | 0.031 | 0.002 |
| P0619_N | 0.011 | 0.018 | 0.060 | 0.021 | 0.000 | 0.000 |
| P0619_P | 0.101 | 0.105 | 0.072 | 0.000 | 0.018 | 0.080 |
| P0619_T | 0.003 | 0.003 | 0.145 | 0.222 | 0.003 | 0.000 |
| P0706_P | 0.010 | 0.058 | 0.010 | 0.000 | 0.000 | 0.019 |
| P0706_T | 0.028 | 0.051 | 0.252 | 0.126 | 0.008 | 0.002 |
| P0729_N | 0.003 | 0.000 | 0.016 | 0.006 | 0.003 | 0.013 |
| P0729_P | 0.020 | 0.081 | 0.054 | 0.000 | 0.000 | 0.071 |
| P0729_T | 0.000 | 0.004 | 0.033 | 0.050 | 0.017 | 0.012 |
| P0913_N | 0.049 | 0.038 | 0.000 | 0.011 | 0.000 | 0.011 |
| P0913_P | 0.084 | 0.158 | 0.134 | 0.000 | 0.000 | 0.039 |
| P0913_T | 0.029 | 0.006 | 0.243 | 0.049 | 0.006 | 0.004 |
| P1010_N | 0.005 | 0.028 | 0.133 | 0.000 | 0.000 | 0.005 |
| P1010_P | 0.049 | 0.171 | 0.046 | 0.000 | 0.000 | 0.003 |
| P1010_T | 0.000 | 0.008 | 0.299 | 0.053 | 0.002 | 0.000 |
| P1011_N | 0.000 | 0.000 | 0.000 | 0.231 | 0.000 | 0.000 |
| P1011_P | 0.202 | 0.154 | 0.123 | 0.003 | 0.003 | 0.084 |
| P1011_T | 0.000 | 0.004 | 0.169 | 0.127 | 0.002 | 0.000 |
| P1118_N | 0.005 | 0.047 | 0.026 | 0.005 | 0.000 | 0.000 |
| P1118_P | 0.133 | 0.254 | 0.023 | 0.000 | 0.000 | 0.152 |
| P1118_T | 0.007 | 0.028 | 0.113 | 0.035 | 0.007 | 0.000 |
| P1120_N | 0.000 | 0.000 | 0.000 | 0.073 | 0.049 | 0.000 |
| P1120_P | 0.124 | 0.084 | 0.084 | 0.000 | 0.026 | 0.061 |
| P1120_T | 0.001 | 0.006 | 0.201 | 0.128 | 0.011 | 0.001 |
| P1202_N | 0.004 | 0.015 | 0.045 | 0.015 | 0.023 | 0.004 |
| P1202_P | 0.121 | 0.088 | 0.058 | 0.000 | 0.006 | 0.000 |
| P1202_T | 0.007 | 0.024 | 0.164 | 0.073 | 0.004 | 0.011 |
| P1208_P | 0.164 | 0.085 | 0.141 | 0.000 | 0.009 | 0.141 |
| P1208_T | 0.005 | 0.014 | 0.153 | 0.032 | 0.014 | 0.000 |
| P1219_N | 0.070 | 0.000 | 0.000 | 0.000 | 0.009 | 0.000 |
| P1219_P | 0.131 | 0.049 | 0.087 | 0.000 | 0.038 | 0.038 |
| P1219_T | 0.000 | 0.000 | 0.055 | 0.008 | 0.016 | 0.000 |

**Table S7.** The real immune cell fraction calculated from single-cell barcode of GSE72056.

| **Sample** | **Bcell** | **Macrophage** | **Endothelial** | **NK** | **CD4 T** | **CD8 T** |
| --- | --- | --- | --- | --- | --- | --- |
| Patient53 | 0.050 | 0.066 | 0.066 | 0.061 | 0.189 | 0.112 |
| Patient58 | 0.024 | 0.012 | 0.000 | 0.024 | 0.116 | 0.387 |
| Patient59 | 0.018 | 0.009 | 0.000 | 0.000 | 0.000 | 0.000 |
| Patient60 | 0.372 | 0.019 | 0.000 | 0.041 | 0.130 | 0.126 |
| Patient65 | 0.052 | 0.021 | 0.000 | 0.000 | 0.096 | 0.344 |
| Patient67 | 0.159 | 0.000 | 0.008 | 0.008 | 0.218 | 0.194 |
| Patient71 | 0.023 | 0.015 | 0.000 | 0.000 | 0.036 | 0.145 |
| Patient72 | 0.168 | 0.000 | 0.005 | 0.005 | 0.291 | 0.103 |
| Patient74 | 0.079 | 0.028 | 0.000 | 0.006 | 0.041 | 0.415 |
| Patient75 | 0.013 | 0.010 | 0.003 | 0.000 | 0.029 | 0.554 |
| Patient78 | 0.013 | 0.000 | 0.000 | 0.000 | 0.000 | 0.000 |
| Patient79 | 0.088 | 0.000 | 0.003 | 0.001 | 0.038 | 0.186 |
| Patient80 | 0.101 | 0.002 | 0.057 | 0.010 | 0.179 | 0.103 |
| Patient81 | 0.013 | 0.004 | 0.009 | 0.004 | 0.039 | 0.109 |
| Patient82 | 0.019 | 0.039 | 0.010 | 0.029 | 0.122 | 0.069 |
| Patient84 | 0.145 | 0.140 | 0.005 | 0.038 | 0.123 | 0.117 |
| Patient88 | 0.044 | 0.106 | 0.003 | 0.023 | 0.094 | 0.122 |
| Patient89 | 0.215 | 0.051 | 0.006 | 0.002 | 0.049 | 0.230 |
| Patient94 | 0.171 | 0.007 | 0.057 | 0.002 | 0.164 | 0.078 |

